# Functional contributions of quantal and non-quantal hair cell synaptic transmission in the vestibular periphery

**DOI:** 10.1101/2025.08.06.668943

**Authors:** Dyllan Zhou, Zhou Yu, Takashi Kodama, Wesley Schoo, Sascha du Lac, Elisabeth Glowatzki, Soroush G. Sadeghi

## Abstract

Information about head motion and gravity is conveyed to the brain by vestibular nerve afferents which are subdivided by their spontaneous firing properties into regular and irregular subtypes thought to be differentially responsible for vestibulo-ocular vs vestibulo-spinal reflexes. In the vestibular periphery, afferents make glutamatergic synapses with type II hair cells (HCs) in all vertebrates. During the evolutionary transition to land, however, amniotes (reptiles, birds, and mammals) additionally developed type I vestibular HCs in which unique calyceal afferent terminals cover the basolateral walls of one or more HCs, enabling a nonquantal (NQ) form of synaptic transmission. Most afferents receive inputs from both types of HCs, but the roles of type I vs type II HCs in generating vestibular afferent firing patterns and behaviors remains unclear. Using optogenetics in mice (both sexes), we confirm that stimulation of type II HCs drives conventional quantal glutamatergic transmission, whereas type I HC stimulation evokes nonquantal responses. In mice with disrupted glutamatergic quantal transmission, NQ transmission effectively drove afferent responses to a wide range of head movement frequencies, as assessed by both vestibular sensory evoked potentials and the vestibulo-ocular reflex. Although the distribution of afferent discharge regularity was unaffected, loss of glutamatergic transmission impaired detection of gravity as evidenced by abnormal contact righting reflex behavior. These results indicate that nonquantal glutamatergic transmission from type I HCs is sufficient to generate normal afferent firing patterns and dynamic vestibular behaviors and that glutamatergic release from type II HCs is required for the detection of gravity.

**Significance Statement:** The vestibular system enables balance and spatial orientation by detecting head movements and gravity. During vertebrate evolution, amniotes developed a unique form of synaptic communication—non-quantal (NQ) transmission—between type I hair cells and calyceal afferent nerve fibers in vestibular sensors. This study reveals that NQ transmission can robustly drive vestibular responses across a wide stimulus range, even in the absence of conventional glutamate-based (quantal) signaling. Using electrophysiology, optogenetics and behavioral assays, we show that NQ and quantal transmission serve distinct but complementary roles: NQ supports rapid, broad-range signaling, while quantal transmission is critical for precise gravity detection. These findings highlight how evolutionary innovations in synaptic transmission enhance vestibular function, contributing to our understanding of balance disorders and sensory processing.

## Introduction

The vestibular system plays a crucial role in maintaining balance and spatial orientation through the detection of head movements and gravity. Vestibular sensors in the inner ear consist of three semicircular canals that provide information about rotational movements and two otolith organs that encode linear acceleration in the horizontal and vertical planes. Head movements are detected by the deflection of stereocilia on hair cells (HC), which convert mechanical stimuli into neural signals. These signals are then transmitted to the brain via vestibular afferent neurons. Vestibular afferent nerve fibers are classified as either ‘regular’ or ‘irregular’ based on the variability of their resting discharge (Goldberg et al., 1984; Goldberg, 2000). This classification correlates with their response properties to head movements: irregular afferents exhibit high-pass, phasic responses that increase in gain and phase lead with higher frequency stimulation, whereas regular afferents display more band-pass, tonic responses with relatively stable gain and phase across frequencies (Fernández and Goldberg, 1976; Lysakowski et al., 1995; Brichta and Goldberg, 1996; Sadeghi et al., 2007b, 2007a; Yang and Hullar, 2007; Lasker et al., 2008). Both types of afferents typically receive inputs from type I and type II vestibular hair cells (HC) (Fernández et al., 1995a; Lysakowski and Goldberg, 1997, 2008a). Type I HCs synapse with the specialized large calyx terminals of afferents that envelop type I HCs, while type II HCs either synapse onto bouton terminals or directly synapse on the outer surface of calyces (Goldberg et al., 1990; Fernández et al., 1995b; Lysakowski and Goldberg, 1997, 2008b; Goldberg, 2000; Desai et al., 2005a, 2005b; Ahmad et al., 2025). Classic vesicular glutamate release from hair cells drives quantal transmission, resulting in excitatory postsynaptic potentials (Rennie and Streeter, 2006; Songer and Eatock, 2013; Sadeghi et al., 2014; Highstein et al., 2015). The enclosed space between type I HCs and calyces facilitates glutamate accumulation and spillover, generating slower synaptic responses (Sadeghi et al., 2014). Calyx terminals also receive unusual non-quantal (NQ) inputs from type I HCs (Yamashita and Ohmori, 1990; Goldberg, 1991; Holt et al., 2007a; Lim et al., 2011; Contini et al., 2012; Songer and Eatock, 2013; Contini et al., 2022, 2024), which result in faster responses than quantal transmission and are mediated by ephaptic transmission by K^+^ and resistive coupling (Contini et al., 2022, 2024; Govindaraju et al., 2023).

The functional significance of the individual components of this complex synaptic network in the vestibular periphery is not yet completely understood. In this study, we examined how quantal and NQ transmission contributed to afferent signaling. Using optogenetic stimulation in 3–4 week old mice, we activated all HCs in the excised vestibular crista and measured the afferent response which included quantal and NQ components. Secondly, we selectively activated type I or type II HCs in the semicircular canal cristae and found that afferent nerve fibers received NQ input from type I HCs and quantal input from type II HCs. To test the role of quantal versus NQ transmission in afferent responses, we blocked glutamatergic quantal signaling using either vesicular glutamate transporter 3 (vglut3) KO mice or intratympanic injection of NBQX. In both cases, vestibular sensory evoked potentials (VsEPs) remained normal, indicating NQ transmission can generate synchronized afferent responses during fast head movements. At the behavioral level, in the absence of quantal transmission the vestibulo-ocular reflex (VOR) remained intact for head movements up to 1 Hz, suggesting that NQ input alone was sufficient for encoding such head movements. Single unit recordings confirmed the presence of regular afferents in vglut3 mice, consistent with a suggested role for these afferents in transmitting vestibular signals that drive the VOR (Minor and Goldberg, 1991). However, mice lacking quantal transmission showed abnormal contact righting reflex, indicating impaired gravity sensing. Thus, NQ transmission is sufficient for encoding dynamic head movements, while quantal transmission is required for detecting gravity.

## Methods

All animals were handled in accordance with animal protocols approved by the Johns Hopkins University Animal Care and Use Committee. Mice of either sex were used at postnatal age of 22 to 30 days (P22-P30) for *in vitro* electrophysiology recordings and immunohistochemistry experiments and at 2 to 5 months of age were used for *in vivo* electrophysiology and behavioral tests. Wild-type (WT) C57BL/6J mice (Jackson Laboratories (JAX) #000664) were used for control experiments. To specifically drive reporter gene expression in vestibular HCs, we obtained Gfi1-Cre mice1 (generously provided by Dr. Lin Gan and Dr. Jian Zuo) (Yang et al., 2010). To selectively activate HCs, we also obtained vglut3-Cre mice (from JAX, Tg(Slc17a8-icre)1Edw/SealJ, JAX#018147) which exhibit sparse and random expression of reporter genes in HCs. To examine the cre expression pattern in a mouse line, they were crossed with Ai3 mice (from JAX, B6.Cg-Gt(Rosa)26Sortm3(CAG-EYFP)/Hze/J; JAX #007903) (Madisen et al., 2010) (Figs. 1B and 3A). To drive the expression of ChR2, Gfi1-Cre or vglut3-Cre mice were crossed with Ai32 mice (from JAX, B6;129S-Gt(ROSA)26Sortm32(CAG-COP4*H134R/EYFP)Hze/J; JAX#012569) (Madisen et al., 2010) (Figs. 2B, 3B1, and 3C1). To eliminate glutamatergic quantal synaptic transmission, vglut3 knockout mice were obtained (from JAX, B6;129S2-Slc17a8tm1Edw/J, JAX# 016931) (Grimes et al., 2011). As described previously (Seal et al., 2008), these mice were deaf and showed no startle reflex to sound.

**Figure 1.**
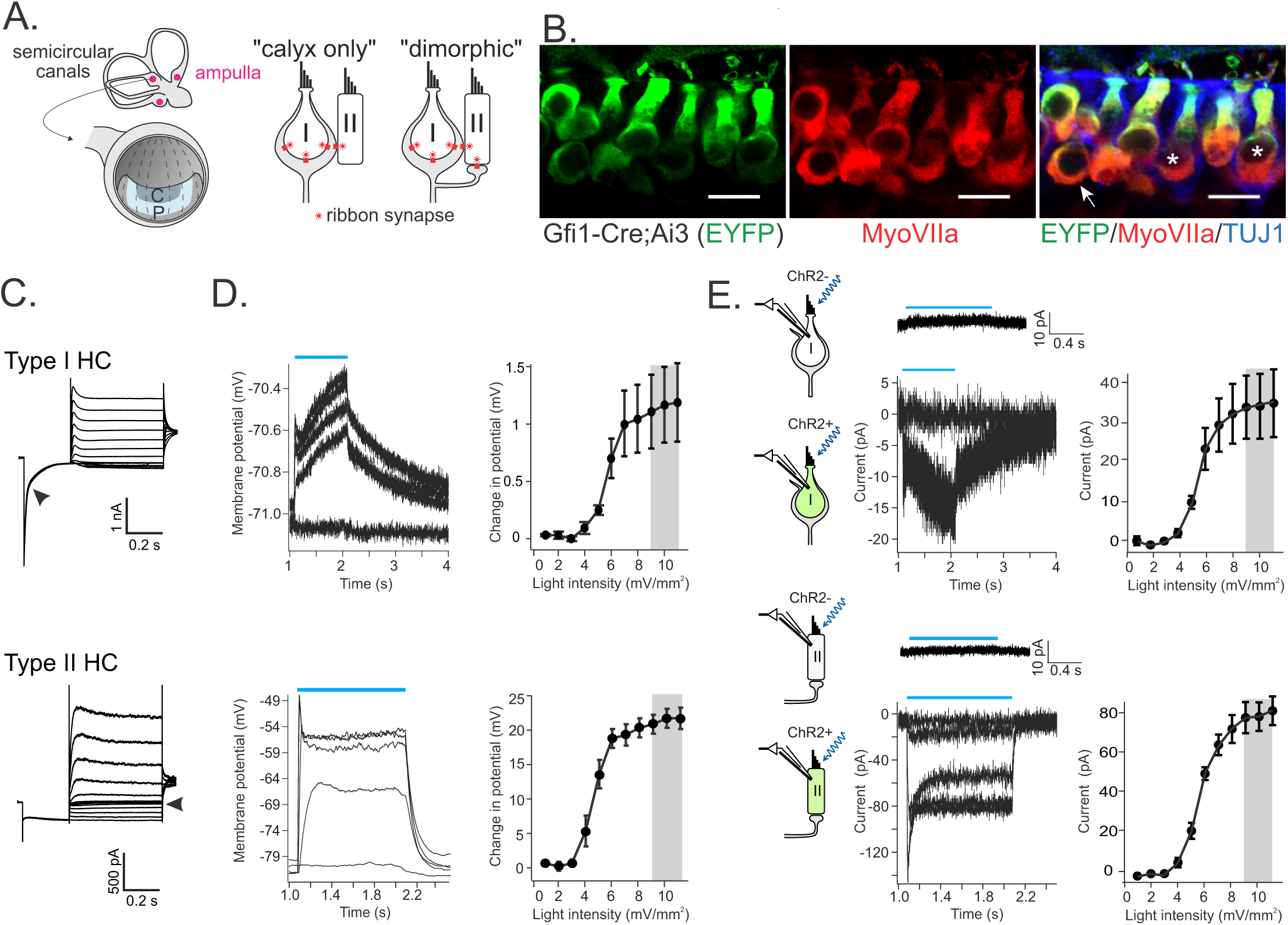
Activation of ChR2 in type I and type II HCs. **A.** Left: peripheral vestibular organs and the neuroepithelium (crista) of canal organs that is divided into central (c) and peripheral (p) zones. Right: schematic of calyx-afferents including both calyx-only and dimorphic types. Note the inputs from type II HCs onto the outer surface of calyces. **B.** In Gfi1-Cre;Ai3 mice, all HCs were EYFP^+^ including type I HCs (asterisks) that were surrounded by calyx afferent terminals and type II HCs (arrow). HCs were immunolabeled by anti MyosinVIIa antibody and afferent fibers by Tubulin J 1 antibody. Scale bar: 10 μm. **C.** Example currents in response to the voltage clamp step protocol. Type I HC shows a typical slow inactivating current during the initial hyperpolarizing step (arrowhead), which is absent in type II HC responses. Type II HC shows high resistance near its resting membrane potential (arrowhead). **D.** Membrane potential of type I HCs and type II HCs were depolarized by light pulses (blue bars) at increasing intensities of 0, 4, 5, 7, and 11 mW/mm^2^. Near-saturating light intensities (right, grey bars) were used to characterize HC to calyx-afferent synaptic transmission. **E.** In the absence of ChR2 expression (control experiments), HCs were not affected by light pulses (blue bars). Example trace was averaged from 10 recordings at a holding potential of −90 mV and light intensity of 9 mW/mm^2^. When ChR2 was expressed, an inward current was induced in type I HCs and type II HCs by light pulses at increasing intensities (same as in D). Note the smaller size of the average currents in type I HCs compared to type II HCs.

**Figure 2.**
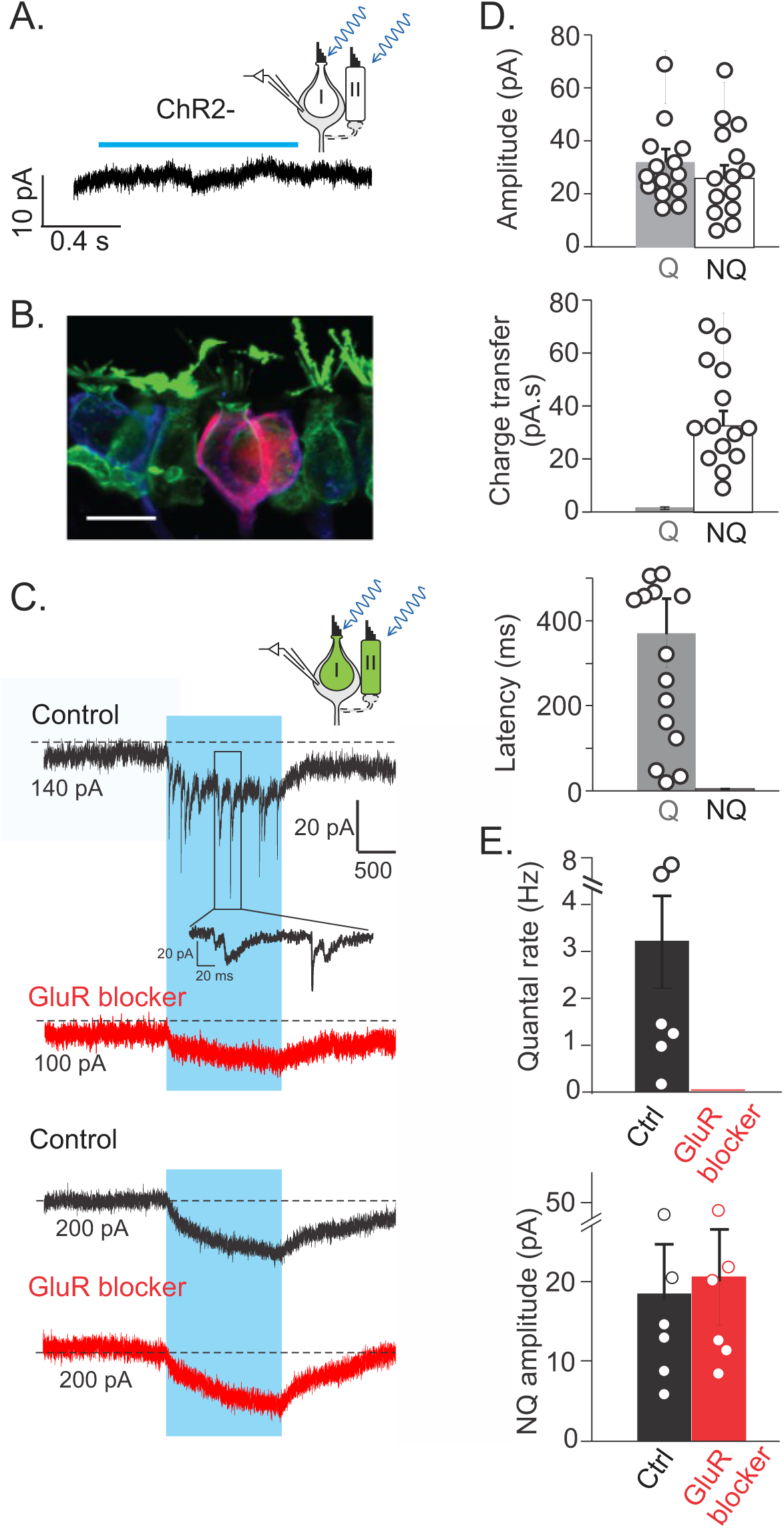
ChR2 activation of HCs results in both quantal and non-quantal (NQ) signals in calyx terminals. **A.** Example voltage clamp recording from a calyx terminal showing that in the absence of ChR2 expression, light pulses (blue bar) did not have any effect on recorded currents (holding potential: −90 mV). **B.** In Gfi1-Cre;Ai32 mice, a recorded calyx-afferent is filled with biocytin (red) and is surrounded by ChR2-YFP positive HCs (green). **C.** Representative responses of calyx terminals during HC activation. When HCs expressed ChR2, optical stimulation resulted in responses with synaptic events (enlarged inset) and a standing NQ component. The quantal component could be inhibited completely by a cocktail of glutamate receptor blockers (GluR blocker, red trace), leaving only the NQ component. Recording on the bottom shows a rare instance with only a NQ response and no quantal component. The NQ response was not affected by GluR blocker cocktail. **D.** Average amplitude, charge (area under the curve), and onset time of NQ responses and the first quantal event (n=14). While quantal and NQ responses have similar amplitudes, NQ responses provide larger charge transfer and have a shorter latency. **E.** With application of GluR blocker, the quantal rate drops to zero, but the NQ amplitude does not change. Note, all currents are from whole-cell recordings voltage clamped at −90 mV.

**Figure 3.**
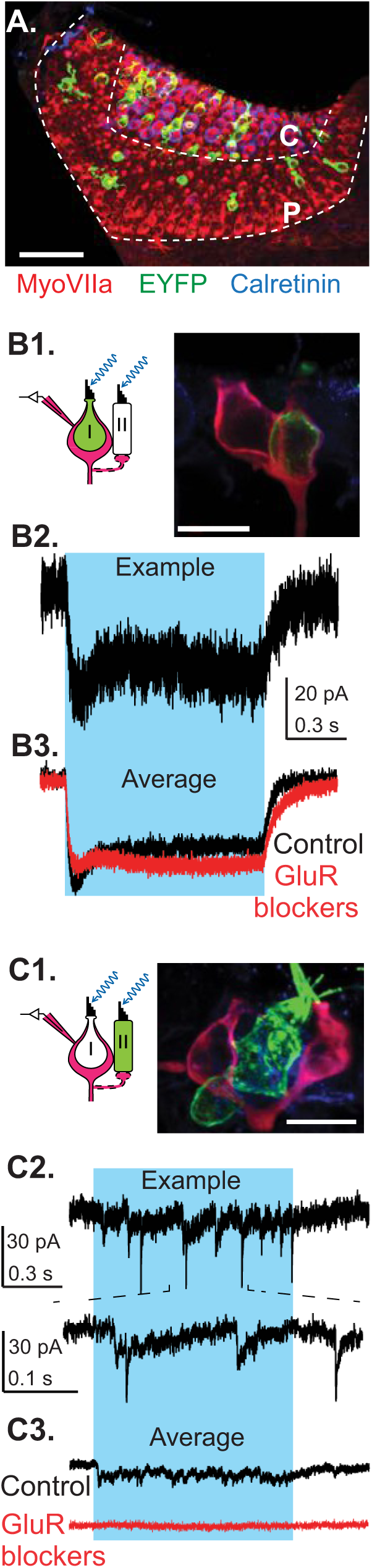
Type I and type II hair cells contribute to quantal and NQ signals, respectively. **A.** Reporter gene expression driven by vgluT3-Cre in the crista (cross between VgluT3-Cre and Ai3 mice). HCs were immunolabeled by MyosinVIIa. Note EYFP^+^ HCs that are sparsely and randomly distributed in both central and peripheral zones of the crista. The central zone was marked by calyx-only afferents immunolabeled with calretinin (Scale bar: 50 μm). **B.** A biocytin-filled recorded afferent contacted one EYFP^+^ type I HC, with no EYFP^+^ type II HC in its vicinity. Representative traces of NQ currents in afferent terminal during ChR2 stimulation of the type I HC are shown below (top: an individual trace; bottom: averaged responses from 10 trials). These NQ responses were not affected by GluR blockers (in blockers vs. Ctrl: −17.1 ± 3.5 pA vs. −18.6 ± 2.6 pA, n=11, p= 0.32, *paired t-test*.). **C**. A biocytin-filled recorded afferent close to an EYFP^+^ type II HC without any EYFP^+^ type I HC nearby. ChR2 stimulation of type II HC generated mainly quantal currents (top: an individual trace and enlarged inset). A standing current was observed in averaged responses from 10 trials. The amplitude of these steady currents correlated with the rate of quantal events (R = 0.891, p = 0.017, n = 6) and were eliminated by application of GluR blocker cocktail, suggesting the quantal glutamate release as their source. All recordings were voltage clamped at −90 mV.

### Tissue preparation

This procedure was performed as previously described (Sadeghi et al., 2014; Ramakrishna and Sadeghi, 2020; Yu et al., 2020; Ramakrishna et al., 2021). In brief, mice were deeply anesthetized by isoflurane inhalation and decapitated. The inner ear tissue was removed from the temporal bone and placed into extracellular solution. The bony labyrinth was opened and part of the membranous labyrinth was dissected, including ampullae of horizontal and superior (anterior) canals and Scarpa’s ganglion. The membranous labyrinth was then opened above the cristae and utricle and remaining cupulae located on top of hair cells were removed to expose the neuroepithelia of the cristae.

### Electrophysiology recording

The preparation was secured on a coverslip under a pin, transferred to the recording chamber, and perfused with extracellular solution at a rate of 1.5-3 ml/min. The extracellular solution contained (in mM): 5.8 KCl, 144 NaCl, 0.9 MgCl2, 1.3 CaCl2, 0.7 NaH2PO4, 5.6 glucose, 10 HEPES, 300 mOsm, pH 7.4 (NaOH). In some experiments, to increase the extracellular K+ concentration up to 40 mM, equimolar NaCl was replaced with KCl. To record at physiological temperature, the external solution was heated through an inline heater (ThermoClamp-1) in a subset of experiments. To perform patch-clamp recording, tissue was visualized with a 40X water-immersion objective, differential interference contrast (DIC) optics (Axioskop2 microscope, Zeiss) and viewed on a monitor via a video camera (Dage MTI LSC 70 or IR1000). For targeted recording of fluorescent structures, a wide-field excitation lamp (X-Cite 120) was used as light source. Patch-clamp recording pipettes were fabricated from borosilicate glass pipettes with 1 mm inner diameters (WPI). Pipettes were pulled with a multistep horizontal puller (Sutter), fire polished, and coated with Sylgard (Dow Corning). Pipette resistances were 5-8 MΩ. The intracellular solution for HC and calyx–afferent recordings contained (in mM): 20 KCl, 110 K-methanesulfonate, 0.1 CaCl2, 5 EGTA, 5 HEPES, 5 Na2 phosphocreatine, 4 MgATP, 0.3 Tris-GTP, 290 mOsm, pH 7.2 (KOH). Cocktail of glutamate receptor blockers consisted of 25 μM NBQX, 25 μM CNQX, 25 μM CPP and 500 nM MCPG and was bath applied. All blockers were purchased from Tocris Bioscience. External solution containing 25 mM or 40 mM K^+^ was bath or focally applied. Focal application of solutions was performed using a gravity-driven flow pipette (∼100 μm in diameter) placed near the recorded calyx, connected to a VC-6 channel valve controller (Warner Instruments, Hamden, CT). CsCl based internal solution was used in experiments in which calyx terminals were voltage clamped in external solution with high concentration of potassium, containing 135 CsCl, 3.5 MgCl2, 0.1 CaCl2, 5EGTA, 5 HEPES, 2.5 Na2ATP (280 mOsm, pH7.2 (CsOH)) and 5 uM QX-314 to block voltage-dependent sodium channels intracellularly. To reconstruct the morphology of recorded cells, 0.25-0.3% biocytin (Sigma) was added to the internal solution. Whole-cell patch-clamp recordings were performed from HCs and calyces in the cristae of superior (anterior) or horizontal canals. All measurements were acquired using pCLAMP10.2 software in conjunction with a Multiclamp 700B amplifier (Molecular Devices), digitized at 50 kHz with a Digidata 1440A, and filtered at 10 kHz. For whole-cell recordings, a voltage clamp protocol that contained a 50 ms long, 10 mV hyperpolarization step from −75 or −80 mV was applied to examine membrane resistance (Rm) and access resistance (Ra). Only recordings that had Ra smaller than 25 MΩ were included in data analysis.

### Cell identification

Putative type I HCs were identified with DIC optics by their narrow necks near the top of the HC and by the calyx nerve terminal around them, which could be visualized as a thickening around basolateral walls of the HC. Type I HCs were accessed for whole-cell recording by separating the surrounding calyx terminals with positive pressure from the recording electrode applied to the top of the HC-calyx synaptic space. Type II HCs were identified by their cylindrical shape and the lack of a calyx terminal. The profile of voltage dependent conductances was examined for every morphologically identified HC by applying a voltage-clamp protocol consisting of a hyperpolarization step from −79 mV to −129 mV followed by depolarization steps to different potentials from −129 mV to −9 mV. For type I HCs, the hyperpolarizing step triggered large, slowly inactivating currents of several nAs, resembling I_K,L_ mediated by delayed rectifier potassium channels (Martin et al., 2024). In contrast, the hyperpolarizing step in type II HCs activated inward currents of a few hundred pAs that were likely mediated by a class of delayed rectifier currents and hyperpolarization activated Ih currents (Brichta et al., 2002; Yu et al., 2020; Meredith et al., 2023; Mohamed et al., 2024).

Calyx terminals were identified under DIC optics as a thickening around type I HCs that continued at the base as a nerve fiber (Songer and Eatock, 2013; Sadeghi et al., 2014; Ramakrishna and Sadeghi, 2020; Ramakrishna et al., 2021). Recordings were mostly performed from the base of calyx terminals. For whole-cell recordings, the same voltage-clamp protocol as used for HCs was also applied to calyces, in which the depolarization step could evoke large inward transients likely mediated by sodium channels (Songer and Eatock, 2013; Sadeghi et al., 2014; Meredith and Rennie, 2020; Contini et al., 2024). The majority of recordings were performed from the central zone of the cristae, as it was more accessible in our preparation. Nevertheless, we also assessed responses from calyx terminals in the peripheral zone, as confirmed by immunostaining against calretinin, which labels calyx-only terminals in the central zone. In Gfi1-Cre; Ai32 mice, all peripheral zone calyx terminals tested exhibited both quantal and NQ synaptic currents during ChR2 activation of HCs, with properties comparable to those obtained from calyx terminals recorded from the central zone. Thus, for our analysis, no qualitative differences in synaptic signals were observed for calyx terminals at the central and peripheral zones.

### Optogenetic stimulation

The intensity and timing of blue light pulses were controlled by a TTL pulse generator (Mightex) and were delivered through an optical fiber (diameter: 910μm, Thorlab) coupled to an LED light source (Xlamp, 485nm, Cree). The optical fiber was placed close to the preparation and recording area. Stimuli consisted of 1 s light pulses at increasing light intensities (0-11 mW/mm^2^) to determine light intensities for getting near saturation responses in HCs, so that the amplitude of synaptic responses in calyces was between 85-95% of the maximal value.

### *In vitro* electrophysiology data analysis

For each HC and calyx terminal recorded in whole cell patch clamp mode the onset, amplitude, 95% rise time (only for HCs), 5% decay time, total charge transfer (only for calyx-afferents), and NQ responses evoked at near saturating level were analyzed from averaged responses of at least 10 trials. The threshold for detecting onset was set as mean plus three standard deviations. The amplitude was calculated as the mean of the response at 500 – 550 ms near the middle of stimulation. Trial to trial variation of response amplitude or total charge transfer was calculated from each trial for at least ten trials in total. Spikes were automatically detected with a threshold set manually. The above analysis was performed in MATLAB (Mathworks, Inc). Quantal synaptic events were analyzed using Minianalysis (Synaptosoft). All synaptic events were detected manually. Parameters of these events were calculated by built-in routines in Minianalysis, and at least 5 points were averaged for a peak value. Decay phase of synaptic events was fitted with single exponential and only fitting results with coefficient of determination R^2^ > 0.85 was considered as a good fit and included for further analysis. For those well fitted events, 10-90% τ_decay_ was calculated. Data are reported as mean ± SE, unless specified otherwise.

### Immunohistochemistry

Freshly excised temporal bone tissue with bony labyrinth opened or tissue after electrophysiology recording was dropped in 4% paraformaldehyde (PFA) for fixation of 1hr to 24 hrs at 4°C and then rinsed in PBS solution. For antibody-based immunolabeling, samples were first incubated in blocking buffer (PBS with 10% normal donkey serum, 0.3% Triton X-100) for 1 hr. Samples were then incubated in primary antibody diluted in blocking buffer for 24-48 hrs at 4°C. Samples were rinsed in PBS before incubation in secondary antibody diluted 1:500 – 1:750 in blocking buffer for 2 hrs. After rinsing in PBS for 3-5 times, samples were mounted on glass slides in FluorSaveTM Reagent medium (EMD Millipore). Primary antibodies and dilutions used in this work were: goat anti-GFP (1:5000, SicGen), mouse anti-TUJ1(1:250, BioLegend, labeling afferents), mouse anti-MyosinVIIa (1:250, Sigma, labeling HCs), rabbit anti-MyosinVI (1:250, Sigma, labeling HCs), rabbit anti-TUJ1(1:250, BioLegend), guinea pig anti-vglut3 (1:500, originally from Dr. Robert Edwards, provided by Dr. Omar Akil). Secondary antibodies used in this work were: Alexa Fluor® (488, 555, or 595) conjugated donkey anti-goat IgG(H+L) (ThermoFisher), Alexa Fluor®(488, 568 or 647) conjugated donkey anti-rabbit IgG(H+L) (ThermoFisher), Alexa Fluor®(488, 568, or 647) conjugated donkey anti-mouse IgG(H+L) (ThermoFisher), Alexa Fluor®594 AffiniPure donkey anti-guinea pig IgG(H+L) (Jackson ImmunoResearch), and Rhodamine Red TM-X(RRX) donkey anti-guinea pig IgG(H+L) (Jackson ImmunoResarch). For labeling biocytin filled HCs and vestibular afferents, tissue was stained by incubating with Alexa Fluor® 488 or 568 conjugated strepdavidin (ThermoFisher) for 2hrs. Fluorescence images were acquired using a laser scanning confocal microscope (LSM700, Zeiss) under the software control of ZEN. Imaged Z-stacks were collected under near saturating laser intensities for each channel.

### Single unit extracellular recording from the vestibular nerve

We used methods described in detail previously (Raghu et al., 2019). Briefly, mice were deeply anesthetized by intraperitoneal injection of ketamine (80 mg/kg) and xylazine (10 mg/kg), were held in a stereotaxic frame, and a craniotomy was performed in the lateral part of the parietal bone. The lateral portion of the cerebellum was aspirated to expose Scarpa’s ganglion and the eighth nerve. Glass electrodes with impedances of 20 – 60 MΩ were filled with 3 M NaCl solution and positioned over the nerve under direct visualization through a surgical microscope (Leica). The electrode was moved up to 300 – 350 μm inside the nerve, using a microdrive (MO-10, Narishige) and resting discharge of afferents were recorded for at least 60 s. Signals from the nerve were amplified and bandpass filtered between 300 Hz and 1 kHz (EXT-02B amplifier, npi). An external auditory speaker was used to discern the activity of neurons. In addition, the voltage from the microelectrode was digitized with a 16-bit A/D converter at a sampling rate of 100 kHz (micro1401, CED). All signals were then recorded on a PC for offline analysis.

### Bilateral Intratympanic injections

Intratympanic injections (IT) of about 20 μl of AMPA receptor antagonist NBQX (20 mM) were performed in each ear through the tympanic membrane under a surgical microscope, using a plastic tube with a sharp tip (about 27 gauge). Following the injection, mice were kept with the injected ear up for 5 min for the drug to penetrate the inner ear and take effect. The other ear was then injected in the same way. The efficiency of IT injections was tested by the absence of auditory brainstem responses (ABR) after injections.

### Recordings of the vestibular sensory evoked potentials (VsEP) and auditory brainstem responses (ABR)

We used VsEP to measure the vestibular nerve (otolith) function, as described previously (Jones et al., 2002, 2011; Raghu et al., 2020). Briefly, after anesthesizing an animal by an intraperitoneal (IP) injection of a mixture of ketamine (80 mg/kg) and xylazine (10 mg/kg), subcutaneous needle electrodes were placed at the vertex (non-inverting recording electrode) and below the pinna of the right and left ears (inverting and ground targets). The head was then held by a hair clip attached to a plate that moved in the vertical axis by an electromagnetic shaker (ET-132, Labworks Inc.). The head was moved in the up or down direction (naso-occipital translation) with jerks (i.e., change in acceleration) of 2 ms duration and amplitudes of 0.7–2 g/ms (in 3 dB steps). Signals were amplified by a Grass P511 amplifier (gain of 100,000), filtered (0.3–3 KHz) and sampled at 100 K by a micro1401 (CED). Responses were averaged over 100–200 trials in each direction to resolve responses from background activity. The peak-to-peak amplitude of the first positive to negative wave (P1-N1) and the latencies of P1 and N1 were measured to quantify the activity of the vestibular nerve.

The anesthetized mouse was then moved to an acoustic chamber for recording the ABR in response to clicks (10 – 90 dB). The mouse was positioned on its stomach in front of a speaker located at a distance of 10 cm. The peak-to-peak amplitude of the first wave of the average trace from 200 recordings at each amplitude was measured to quantify the response of auditory nerve fibers. Following IT injection of NBQX, a decrease of at least 80% in ABR amplitude in response to 90 dB click was considered as evidence for penetration of the drug into the inner ear. As a control for IT injections, we measured the change in ABR amplitude and threshold after IT injection of extracellular solution. Initially, to verify penetration of NBQX into the inner ear following IT injections and the duration of its effect, ABR and VsEP recordings were performed in a group of mice every 30-60 min for up to two hours (n = 5) and up to 3 hours (n = 2).

### Recording the vestibulo-ocular reflex (VOR)

Eye movement recordings using video-oculography (Stahl et al., 2000) were performed with methods described previously (Kodama and du Lac, 2016). In brief, a headpost was implanted on the skull under isoflurane anesthesia. After a recovery period of 2-5 days, mice were acclimatized to the VOR setup by being head restrained in a tube and rotated in the dark for a few minutes/day at different frequencies for 2-3 days. Rotations at 3 frequencies (0.25, 0.5, and 1 Hz; peak velocity 15.7 deg/s) were used to assess the VOR in WT and vglut3 KO mice. For accurate eye tracking, physostigmine salicylate (0.05 %) or pilocarpine (1 – 2%) eye drops were used to limit pupil dilation. Eye position signals were recorded at 200 Hz with a miniature infrared video camera and differentiated digitally to obtain eye velocity. Rapid transients in eye velocity (>200 deg/s) due to quick phases (saccades) were removed digitally. Gain and phase were assessed by fitting sinusoids to eye velocity data. Analyses were done using Igor Pro.

### Contact righting reflex

To identify whether mice detected a change in the gravity axis, we used the contact righting reflex (Feather-Schussler and Ferguson, 2016; Grin’kina et al., 2016). Mice were lightly anesthetized by isoflurane inhalation and were placed between two surfaces in a supine position. The top surface was transparent so that we could measure the time from when an animal came out of anesthesia (i.e., started moving) to when it flipped over (i.e., at least three limbs were in contact with the bottom surface). The contact righting reflex was assessed in vglut3 KO mice as well as in WT mice prior to and after bilateral IT injection of NBQX.

### Experimental design and statistical analysis

Experimenters were blinded to genotype during data acquisition for VOR and VsEP recordings from WT vs. vglut3 KO mice. To avoid effects of age on results, animals of comparable ages were used for all experiments across different approaches (e.g., VsEP in mice that received NBQX injection and vglut3 KO mice). Statistical significances of comparisons between values obtained from HCs/calyces before and after a manipulation were assessed by paired two-tailed Student’s t-test. Results for VsEP (amplitudes and latencies for different stimulus accelerations) and VOR (gain and phase for different frequencies of rotation) were compared between conditions and animals using analysis of variance (ANOVA) or repeated measures ANOVA, with post hoc Bonferroni test, to account for multiple comparisons. Some of the analysis was done in MATLAB (MathWorks) and scripts that were custom written for the analyses in this study, based on algorithms that have been used previously (Kodama and du Lac, 2016; Raghu et al., 2019, 2020). Student t-tests were performed in Excel and ANOVA was performed using PRISM (GraphPad).

### Code and data accessibility

The analysis codes and data will be made available upon reasonable request.

## Results

### Vestibular hair cell depolarization via optogenetic stimulation

Previous studies have used stimulation of hair bundles to investigate inputs from individual HCs to a single recorded vestibular afferent fiber (Songer and Eatock, 2013; Ono et al., 2024). Here, the aim was to monitor the combined HC synaptic inputs to individually recorded afferents by optogenetically stimulating all HCs in excised vestibular mouse cristae. The expression of Cre in Gfi1-Cre mice in all hair cells was confirmed by crossing them with an Ai3 reporter line (Fig. 1B). To express ChR2 exclusively in HCs, we crossed Gfi1-Cre mice (Yang et al., 2010) with an Ai32 line that carries ChR2 (Madisen et al., 2010) and the expression was confirmed by co-expression of fluorescence in HCs (Fig. 2B). We tested the level of HC depolarization that could be achieved with optogenetic stimulation using HC whole-cell patch-clamp recordings from acutely excised crista tissue of 3-4 week-old mice. Every HC in ChR2-positive tissue could be excited by blue light pulses (n=32), whereas HCs in ChR2-negative preparations did not respond (n=6). Type I and type II HC recordings were identified by their distinctive responses to a voltage step protocol (Fig. 1C). Increasing illumination levels were tested, and at near saturating illumination levels, HC responses had rapid onsets, within 1-2 ms of stimulus onset. One-second-long light pulses depolarized type I HCs by only 1.1 ± 0.3 mV, from −76.9 ± 1.5 mV to −75.8 ± 1.6 mV, and type II HCs ∼20 fold more, by 21.9 ± 1.7 mV, from −74.9 ± 2.4 mV to −53.0 ± 1.2 mV (n=7, for each HC type) (Fig. 1D). Hair bundle displacements have also been shown to cause much smaller depolarizations of type I versus type II HCs, which has been attributed to the low input resistance of type I HCs (62.3 ± 5.9 vs. 663.5 ± 50.5 MΩ for type II HCs) caused by an active potassium conductance at resting membrane potential (Martin et al., 2024). However, compared to type II HCs, type I HCs also showed a smaller inward current with optical stimulation (20.1 ± 3.7 pA vs. 67.7 ± 9.6 pA) (Fig. 1E), suggesting a lower level of ChR2 function in type I HCs. Note for the interpretation of data below, the small optogenetic depolarization in type I HCs is unlikely to trigger significant calcium influx to drive glutamate release at type I HC ribbon synapses (Bao et al., 2003). The combination of a small depolarization in type I HCs and a larger depolarization in type II HCs, mostly due to their difference in input resistance, could represent their relative activation levels *in vivo* in response to stimulation.

### Quantal and non-quantal inputs to afferent fibers

We next investigated afferent responses to optogenetic stimulation of all HCs and the contribution of quantal and NQ components to these responses. Most afferent fiber recordings were performed from central zones of the cristae (Fig. 1A), which are innervated by irregular afferents (Lysakowski et al., 1995, 1995; Goldberg, 2000; Eatock and Songer, 2011). The patch pipette specifically approached the calyx ending of an afferent fiber for whole cell recording. Note that such recordings will monitor all nearby inputs that the recorded fiber receives, both from one or several type I HCs the calyx is ensheathing, as well as from type II HCs, that may synapse onto the calyx or synapse onto bouton endings that feed into the same fiber as the recorded calyx ending (in dimorphic fibers) (Fig. 1A).

In WT mice and in the absence of ChR2 expression, optical stimulation did not result in any change in the baseline currents recorded from calyces (Fig. 2A). When ChR2 was present in HCs and recordings from calyx terminals were made (Fig. 2B), optogenetic activation of all HCs activated quantal excitatory postsynaptic current (qEPSC) and resulted in an average increase of quantal qEPSC rates from 0.04±0.01/s to 3.35±0.79/s (n=14, p=0.001, paired t-test) (Fig. 2C), with 13 of 14 fibers showing increased EPSC rates. For all experimental conditions combined, all recorded calyx terminals showed steady inward currents with rapid onset (n=27) (Fig. 2C), with waveforms similar to the reported ‘non-quantal’ (NQ) responses induced in calyx terminals by mechano-transduction of immature mouse utricle (Songer and Eatock, 2013) or type I HC depolarization in calyx terminals of mice utricle (Contini et al., 2024), turtle utricle (Contini et al., 2022), and turtle crista (Holt et al., 2007a). Additionally, to verify whether quantal and/or NQ transmission had a zonal preference, we recorded a subpopulation of calyces (n=5) from the peripheral zones of the cristae that are innervated by regular afferents and observed both quantal and NQ responses.

Light-evoked NQ and quantal currents were comparable in peak amplitude (28.7±4.6 pA vs. 31.1±3.8d pA, p=0.60, n=14, *paired t-test*) (Fig. 2D, top), however, NQ currents produced a much larger total charge transfer (35.4±5.1 pC vs. 0.8±0.2 pC, p<0.0001, n=14, *paired t-test*). Additionally, NQ currents activated faster than qEPSCs in response to ChR2 stimulation (5.4±0.7 ms vs. 284.9±49.5 ms, p<0.0001, n =14, *paired t-test*) (Fig. 2D, middle) and more reliably with little trial-to-trial variation. Note that the rise times of NQ currents here are affected by the speed of ChR2 stimulus and do not represent what naturally occurs in response to a mechano-transduction stimulus. Nonetheless, quantal responses were much slower than NQ responses that were almost instantaneous (Fig. 2D, bottom).

Consistent with previous studies (Holt et al., 2007a; Sadeghi et al., 2014; Highstein et al., 2015), qEPSCs were completely blocked by a cocktail of glutamate receptor (GluR) antagonists (AMPA/kainate receptors: 25 μM NBQX, 25 μM CNQX; NMDA receptors: 25 μM CPP; mGluRI/II receptors: 500 nM MCPG; n=6 recordings) (Figs. 2C and 2E). In contrast, the amplitude of the NQ component was not affected (Blockers vs. Ctrl: 20.7 ± 6.0 pA vs. 18.5 ± 6.3 pA, n= 6 recordings, p = 0.25, *paired t-test*) (Figs. 2C and 2E), consistent with former results describing that ephaptic transmission by K^+^ and resistive coupling underlie these synaptic signals (Contini et al., 2022; Govindaraju et al., 2023). In summary, the glutamate receptor blockade allowed isolation of NQ responses pharmacologically.

Notably, the average decay kinetics of qEPSCs (tau = 6.9±0.5 ms, n=14 calyces, 540 qEPSCs) was faster compared to those reported previously (Sadeghi et al., 2014) due to glutamate accumulation and spillover (t-test compared to 18.7 ± 2.8 ms from Sadeghi et al. 2013, n = 46 calyces, 4633 qEPSCs, p = 0.03). The absence of substantial glutamate accumulation suggests that these qEPSCs originated from type II hair cells, which are more efficiently depolarized by optogenetic stimulation than type I hair cells. In support of this idea, the amplitudes of NQ currents were not significantly correlated with qEPSC amplitudes or with the qEPSC rate, suggesting that these two synaptic signals are unlikely to share a common upstream cellular origin. We therefore hypothesized that in our recordings, the major contribution of type I and type II HCs were to NQ and quantal signals, respectively.

### Type I HC stimulation generates NQ responses, whereas type II HC stimulation generates quantal responses

To investigate how type I and type II HC inputs contribute to the afferent response, we used vglut3-Cre;Ai32 mice to selectively excite type I or type II HCs. Although all HCs in the crista are immuno-positive for the vesicular glutamate transporter-3 (vglut3) (see Fig. 4A), the vglut3-Cre drove sparse expression of reporter genes in only about 20% of hair cells (Fig. 3A). Comparable to responses measured in Gfi1-Cre;Ai32 mice, 1 s light stimulation depolarized ChR2-EYFP^+^ type I HCs by 1.8 ± 0.6 mV and ChR2-EYFP^+^ type II HCs by 22.4 ± 2.8 mV in cristae of vglut3-Cre; Ai32 mice (n=4, each). The sparse distribution of ChR2-EYFP^+^ HCs allowed for selective examination of inputs from type I or type II HCs by targeting calyx-afferents for recording in two scenarios that were selected ‘by eye’: (1) ‘type I HC only’ calyx-afferents with at least one calyx-enclosed ChR2-EYFP^+^ type I HC and no nearby ChR2-EYFP^+^ type II HCs (at least 20 μm around the type I HC) (Fig. 3B1) and (2) ‘type II HC only’ calyx-afferents with no calyx-enclosed ChR2-EYFP^+^ type I HCs nearby, but with at least one ChR2-EYFP^+^ type II HC beside the recorded calyx, that could possibly provide input to the recorded calyx (Fig. 3C1).

**Figure 4.**
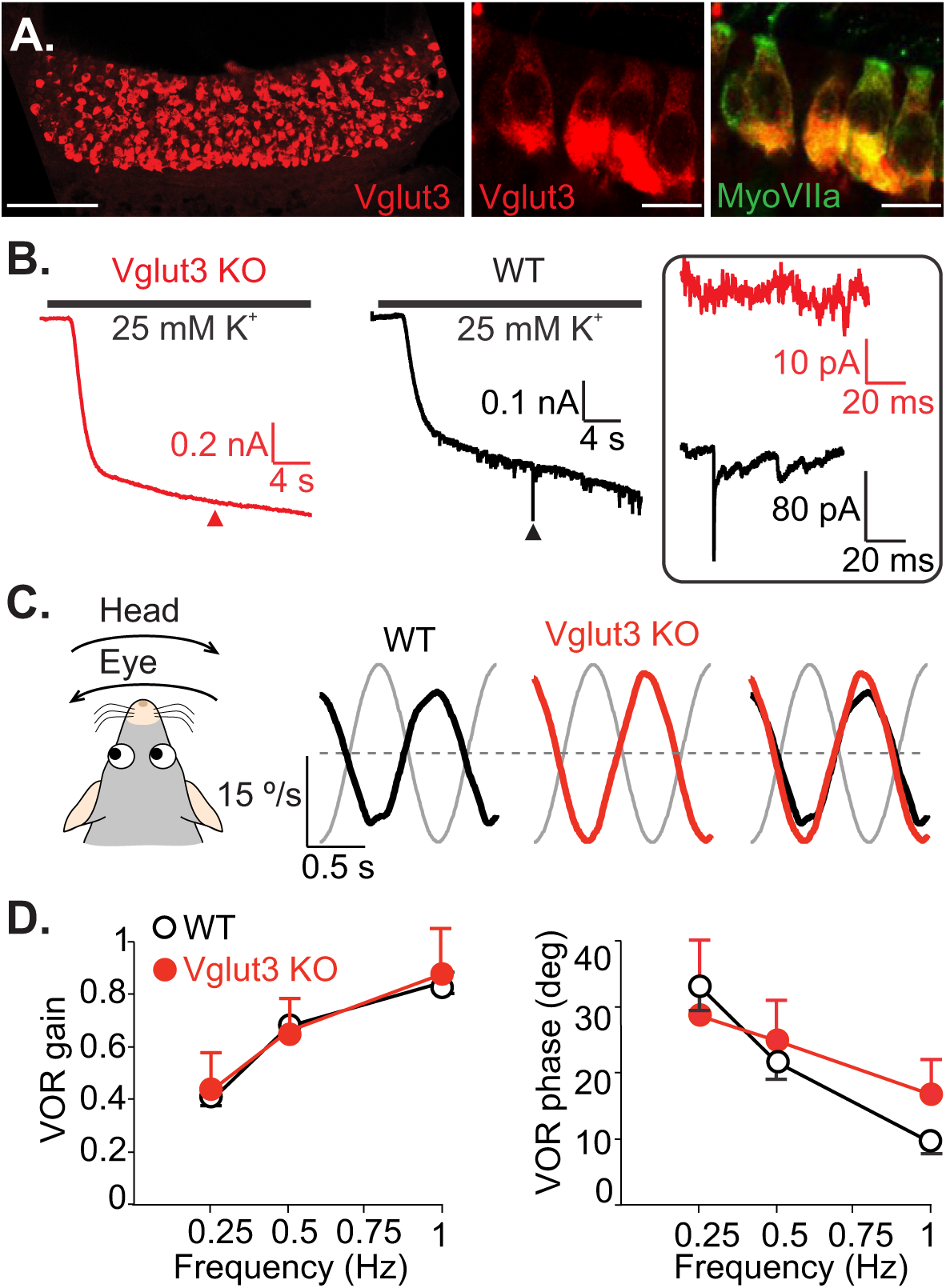
Lack of quantal transmission in vglut3 KO mice does not affect the VOR. **A.** Example of a crista immunolabeled against vglut3 (red); scale bar: 100 μm. Insets on the right show vglut3 (red) and HCs (green, MyosinVIIa antibody). Overlayed image shows that HCs were vgluT3 positive. Scale bars of insets: 10 μm. **B.** Representative inward currents recorded from calyx terminals in WT mice and vglut3 KO mice, at holding potential of −90 mV. Depolarization of HCs by applying external solution with 25 mM [K^+^] increased the frequency of synaptic events in WT mouse, but no synaptic events were observed in vglut3 KO mouse. Currents at arrowheads are shown in inset (box). **C.** Example turntable (head) and eye velocity traces are shown for WT and vglut3 KO mice. Overlap shows the similarity of the response in vglut3 KO mice to that of the WT animal. **D.** Average VOR gain and phase were similar between vglut3 KO (n = 8) and WT (n = 10) mice for all tested frequencies. For frequencies of 0.25, 0.5, and 1 Hz (peak velocity of 15.7 deg/s), VOR gains (mean ± SD) in vglut3 KO mice were 0.44 ± 0.14, 0.65 ± 0.13, 0.88 ± 0.17 and phase leads (mean ± SD) were 28.9 ± 11.20°, 25.1 ± 6.02 °, 16.9 ± 5.34 °, respectively. In WT mice, gains were 0.41 ± 0.04, 0.68 ± 0.03, 0.83 ± 0.03 and phase leads were 33.4 ± 3.87 °, 21.6 ± 2.48 °, 9.82 ± 1.81 °, respectively.

As expected, in all ‘type I HC only’ calyx-afferents tested, ChR2 activation of HCs only induced NQ synaptic currents (19.5 ± 2.5 pA; n=11 at room temperature and n=4 at 32°C; reconstructed morphology for 5 afferents) (Fig. 3B2), similar to NQ responses observed in Gfi1-Cre;Ai32 mice. The NQ responses were not affected by GluR blockers (Fig. 3B3) (Blockers vs. Ctrl, 17.1 ± 3.5 pA vs. 18.6±2.6 pA, n = 11, p = 0.32, paired t-test). No significant increase of qEPSCs was observed (stimulus vs. background, 0.10 ± 0.06/s vs. 0.14±0.06/s, P=0.14, n = 15). About 43 % of ‘type II HC only’ calyx-afferents (n = 6 out of 14 tested) responded to HC activation with an increase of qEPSC rate (stimulus vs. background, 4.77 ± 1.30/s vs. 0.13 ± 0.08/s, n = 6, p = 0.015, 4 afferents with reconstructed morphology) (Fig. 3C2). The unresponsive ‘type II HC only’ calyx-afferents were most likely not connected to the nearby ChR2-EYFP^+^ type II HCs. Notably, qEPSCs induced due to glutamate release from type II HCs, exhibited a range of decay kinetics (Fig 3C2) and when averaged, a small steady inward current was observed during light stimulation (3.3 ± 2.2 pA, n=6) (Fig. 3C3). The latter’s total charge transfer was proportional to the frequency of qEPSCs and was completely abolished by GluR blockers (Fig. 3C3), indicating the presence of some glutamate accumulation even between type II HCs and calyx terminals.

These results indicate that type II HCs supply glutamatergic signals based on quantal release to calyx-afferents. On the other hand, type I HCs were both necessary and sufficient for the presence of NQ synaptic signals. As mentioned previously, the lack of quantal signals from type I HCs is not unexpected, since the small optogenetic depolarization in type I HCs may not trigger significant calcium influx required for vesicular release. However, because of the low membrane resistance of type I HCs, this could well be the case *in vivo*, during head movements, such that type I HCs may transmit signals to the calyx terminal mainly through NQ transmission.

### Non-quantal transmission is sufficient for generating normal VOR responses

To examine the functional role of non-quantal (NQ) inputs, we used vglut3 KO mice, which lack expression of the vesicular glutamate transporter 3. Previous studies have shown that these mice exhibit a complete loss of glutamatergic quantal transmission in the cochlea, resulting in absent auditory brainstem responses (Seal et al., 2008). In wild-type (WT) mice, we observed strong expression of vglut3 in vestibular hair cells (HCs) (Fig. 4A). Depolarizing HCs using high-potassium (25 mM and 40 mM) external solution increased the frequency of spontaneous quantal excitatory postsynaptic currents (qEPSCs) in calyx afferents in WT and heterozygous mice (11/13 calyces, 7 mice). In contrast, vglut3 KO littermates showed neither spontaneous qEPSCs nor any response to high-potassium solution (0/10 calyces, 8 mice) (Fig. 4B). These findings indicate that vglut3 is essential for quantal synaptic transmission from vestibular HCs to afferent terminals, consistent with its known role in cochlear inner HCs and zebrafish lateral line HCs (Obholzer et al., 2008; Seal et al., 2008).

As a first step in assessing the behavioral relevance of quantal versus NQ transmission, we evaluated VOR performance in vglut3 KO mice — a behavioral output that reflects vestibular function. Eye movements were recorded in head restrained mice rotated in darkness across different frequencies (Fig. 4C). Surprisingly, vglut3 KO mice displayed VOR eye movements similar to WT controls, with comparable compensatory responses in the direction opposite to head rotation. The VOR gain did not differ significantly between WT (n = 10) and KO (n = 8) mice at any tested frequency (ANOVA, Frequency × Genotype interaction, F(2, 48) = 0.70, p = 0.50) (Fig. 4D). Although VOR phase at 1 Hz was slightly elevated in KO mice (mean ± SD: WT = 9.8 ± 1.8°, KO = 16.9 ± 5.3°; ANOVA, Frequency × Genotype interaction, F(2, 48) = 4.64, p = 0.01; Bonferroni post hoc, p = 0.04), there were no significant phase differences at 0.25 Hz or 0.5 Hz (Bonferroni post hoc, p = 0.29 and p = 0.45, respectively). These results suggest that NQ transmission from type I HCs alone is sufficient to support VOR behavior in the absence of quantal input.

The VOR is believed to be primarily mediated by regular afferents, which predominantly innervate the peripheral zones of the vestibular neuroepithelium (Minor and Goldberg, 1991). In vglut3 KO mice, we confirmed that quantal transmission was absent in the peripheral crista zone, even when 25 mM K⁺ was applied (n = 5 recordings). We then investigated whether the loss of quantal input in KO mice altered afferent firing patterns. Since it was proposed that afferent discharge regularity is associated with the number of bouton synapses from type II HCs (Huwe et al., 2015; Holmes et al., 2017), a reduction in regular afferents might be expected in vglut3 KO mice, which lack functional type II HC input. However, single-unit extracellular recordings from vestibular afferents in vglut3 KO mice (n = 52 afferents, 3 animals) revealed both regular (65%, n = 34) and irregular (35%, n = 18) firing patterns (Fig. 5A). The coefficient of variation (CV*) for regular afferents was 0.07 ± 0.007 (range: 0.003–0.14), and for irregular afferents was 0.3 ± 0.03 (range: 0.16–0.71), closely matching values reported in WT mice (Raghu et al., 2019) (Fig. 5B).

**Figure 5.**
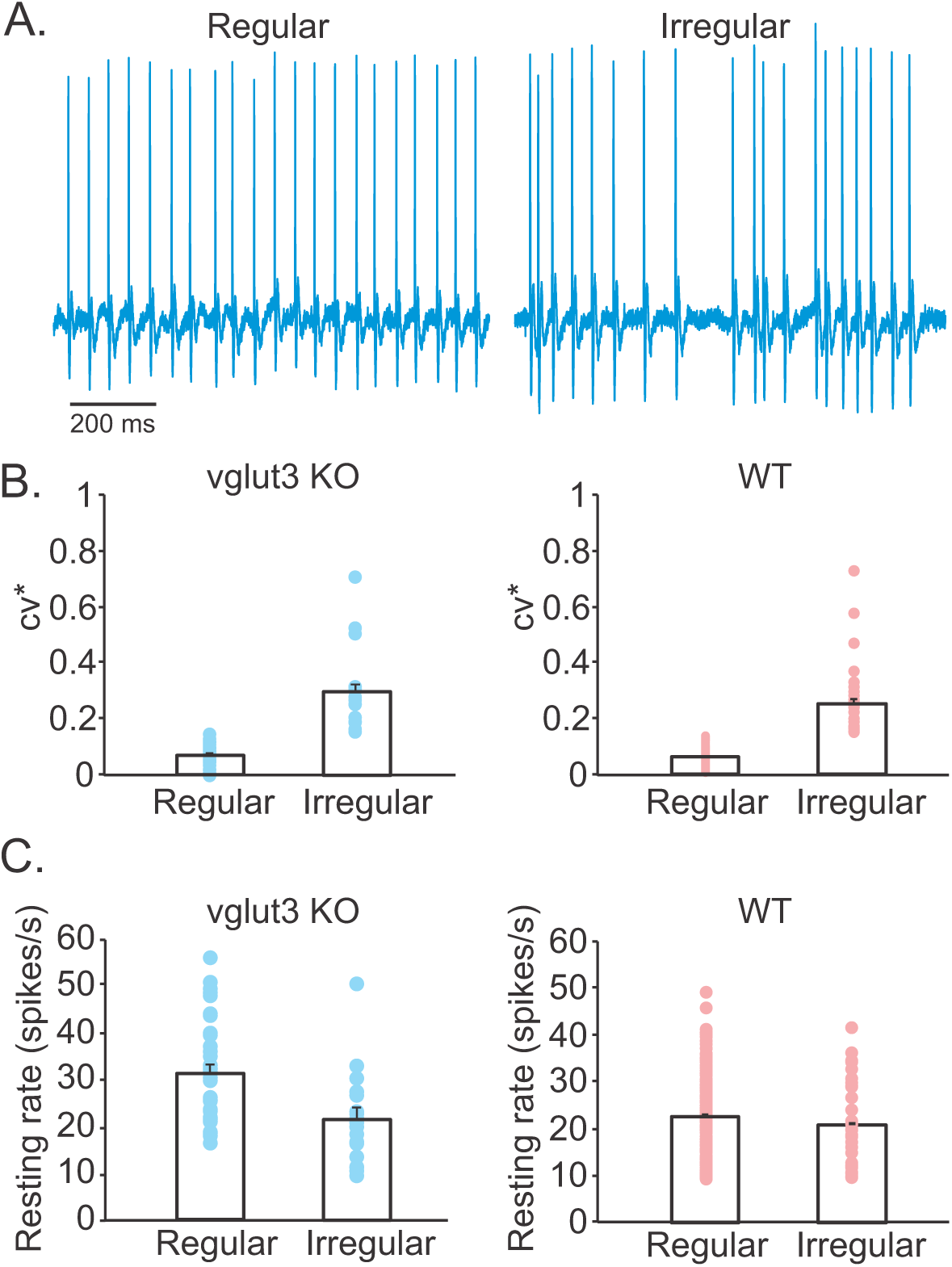
Both regular and irregular afferents are present in the absence of quantal inputs in vglut3 KO mice. **A.** Example extracellular single unit recordings of the resting discharge of a regular and an irregular vestibular nerve fiber from a vglut3 KO mouse. **B.** Averages and individual CV* values calculated for regular and irregular afferents recorded from vglut3 KO (blue, regular: 0.07 ± 0.007, irregular: 0.3 ± 0.03) and WT (red, regular: 0.06 ± 0.001, irregular: 0.25 ± 0.02) mice. **C.** Averages of resting discharges of regular and irregular fibers recorded from vglut3 KO (blue, regular: 31.4 ± 1.9, irregular: 22.2 ± 1.7) and WT (red, regular: 22.8 ± 0.6 spikes/s, irregular: 20.6 ± 1.2 spikes/s) mice. CV* were not calculated for afferents with resting discharges of less than 10 spikes/s and these cells were excluded from the analysis.

Supporting this, a recent *in vitro* study of vestibular ganglion neurons in vglut3 KO mice also identified neuron subtypes with electrophysiological properties similar to those in WT mice (Núñez et al., 2024). Furthermore, the average resting discharge rate of irregular afferents in KO mice (22 ± 2 spikes/s; range: 11–51 spikes/s) was indistinguishable from that in WT animals (Raghu et al. 2019) (21 ± 1 spikes/s; range: 11–41 spikes/s; n = 34; t-test, p = 0.76) (Fig. 5C). Interestingly, the average resting discharge rate of regular afferents in vglut3 KO mice was modestly elevated (31 ± 2 spikes/s; range: 17–56 spikes/s) compared to WT controls (Raghu et al. 2019) (22 ± 1 spikes/s; range: 11–49 spikes/s; n = 173; t-test, p < 0.0001), with a slight rightward shift in the distribution of firing rates.

### Non-quantal transmission is sufficient for normal vestibular nerve responses to rapid transient head movements

To investigate the contribution of NQ inputs to responses of vestibular nerve afferents during accelerative transients of head movements, we recorded field potentials from the vestibular nerve in anesthetized 2-3 month-old mice using established methods for measuring vestibular sensory evoked potentials (VsEP) via high frequency (2 ms duration) linear movements (Jones et al., 2002, 2011; Raghu et al., 2020). We used vglut3 KO mice to determine whether VsEPs are altered in the absence of quantal vesicular release of glutamate from HCs. The absence of glutamatergic transmission in the ears of these mice was further confirmed by the absence of ABR responses to clicks in vglut3 KO mice (n = 5), consistent with previous studies (Seal et al., 2008). In contrast, VsEP responses in the same mice (Fig. 6A) showed amplitudes and latencies similar to WT mice (n = 5, ANOVA, p = 0.1) (Fig. 6B). For the three largest stimuli, average P1N1 amplitudes in WT mice were 0.8 ± 0.04, 1.2 ± 0.08, 1.8 ± 0.14 μV, respectively and in vglut3 KO mice were 1.3 ± 0.2, 1.6 ± 0.2, 2.2 ± 0.3 μV, respectively. For these stimuli, average P1 latencies in WT mice were 2.4 ± 0.05, 2.2 ± 0.02, 2.0 ± 0.01 ms, respectively and in vglut3 KO mice were 2.25 ± 0.06, 2.1 ± 0.05, 2.03 ± 0.01 ms, respectively.

**Figure 6.**
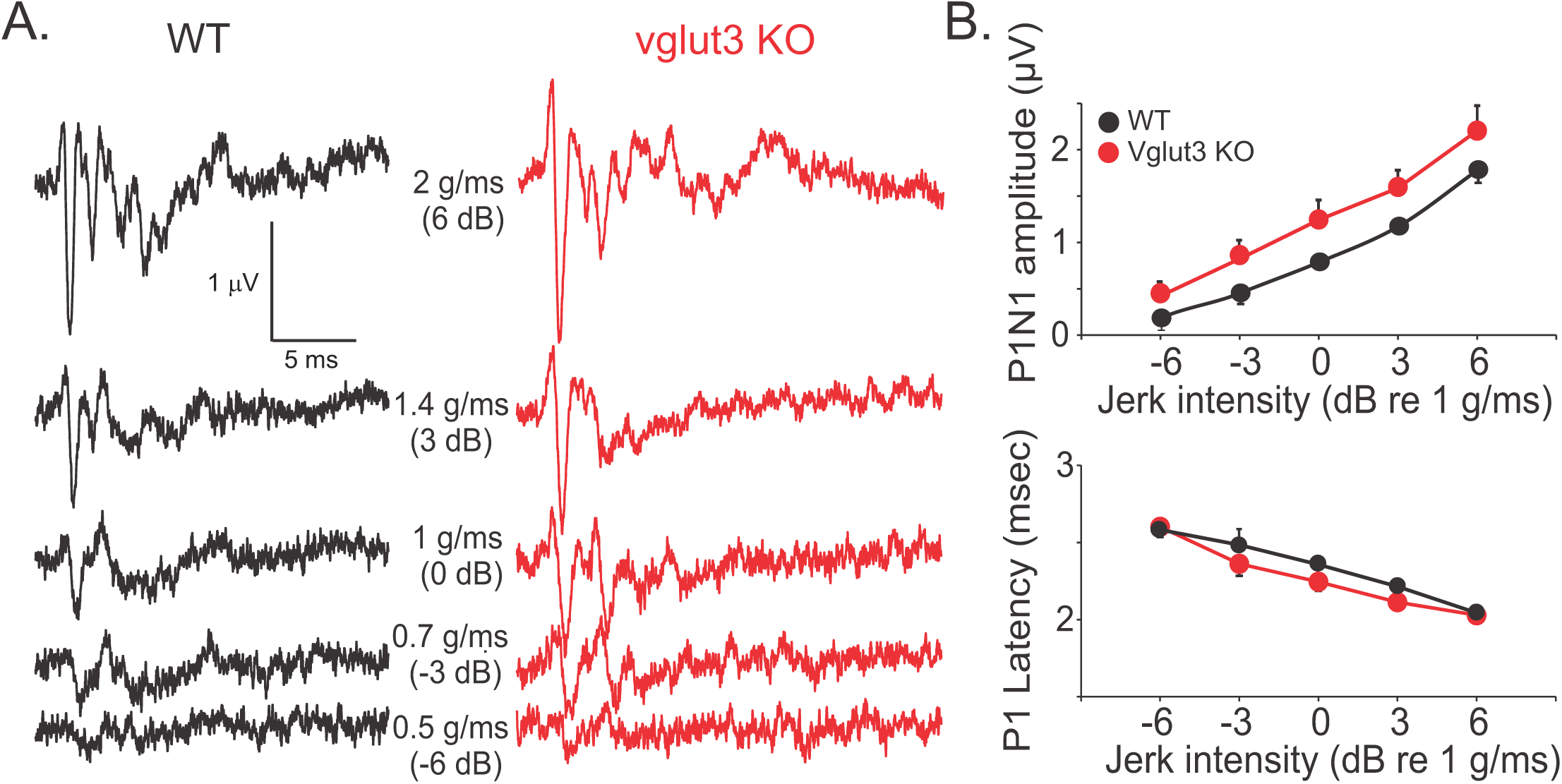
Non-quantal synaptic transmission is sufficient for generating VsEP in vglut3 KO mice. **A.** Example VsEP recordings for a WT mouse and a vglut3 KO mouse showing normal responses in vglut3 KO mouse. **B.** Average amplitudes and latencies of VsEP responses recorded from vglut3 KO mice (n = 5) were similar to WT mice (n = 5, ANOVA, p = 0.1), suggesting that NQ transmission is sufficient for generating VsEP responses. For the three largest stimuli, average P1N1 amplitudes in WT mice were 0.8 ± 0.04, 1.2 ± 0.08, 1.8 ± 0.14 μV and in vglut3 KO mice were 1.3 ± 0.2, 1.6 ± 0.2, 2.2 ± 0.3 μV, respectively. For these stimuli, average P1 latencies in WT mice were 2.4 ± 0.05, 2.2 ± 0.02, 2.0 ± 0.01 ms, respectively and in vglut3 KO mice were 2.25 ± 0.06, 2.1 ± 0.05, 2.03 ± 0.01 ms, respectively.

To further support the above findings in vglut3 KO mice, we used bilateral intratympanic (IT) injection of NBQX, an AMPA receptor antagonist in WT mice to generate an acute model of quantal glutamatergic inhibition. A recent study in guinea pigs showed that application of glutamate blockers on vestibular epithelium in the inner ear did not affect vestibular nerve field potentials (Pastras et al., 2023). We recorded VsEP responses before and after intratympanic (IT) injections and verified that NBQX entered the inner ear after IT injections by recording ABR in response to click stimulation. Since the only known mode of transmission between auditory hair cells and afferent nerve terminals in the cochlea is glutamatergic (Grant et al., 2010, 2011; Martinez-Monedero et al., 2016; Wu et al., 2016; Siebald et al., 2023), ABR wave I amplitudes were expected to be diminished or absent after IT injection of NBQX. We measured wave I amplitudes after IT injection of extracellular fluid as a ‘control’ for the effect of perforation of the tympanic membrane and the presence of fluid in the middle ear on ABR responses (Fig. 7A). In WT mice, ABR and VsEP were measured before and every 30 min after IT injection of NBQX or extracellular solution for up to 3 hours. The amplitude of ABR wave I, which reflects auditory nerve activity, dropped significantly after NBQX injection compared to control values (4.5 ± 0.9 µV, n = 3), reaching 0.6 ± 0.1 µV (n = 5) for a 90 dB stimulus at 30 minutes post-injection. The reduction persisted at 1 hour (Fig.7A) and was still evident at 3 hours (n = 2). While the ABR wave I was significantly decreased during this time, the VsEP recordings did not show any significant change in the peak-to-peak amplitude of the first positive and negative waves (P1N1) or P1 latency (Fig. 7B). This finding supports the idea that VsEP responses represent the synchronized responses of the most phasic striolar afferents with complex calyx terminals that innervate type I HCs (Ono et al., 2020). Together, these findings show that NQ transmission is sufficient for generating afferent responses to rapid transients in head movements.

**Figure 7.**
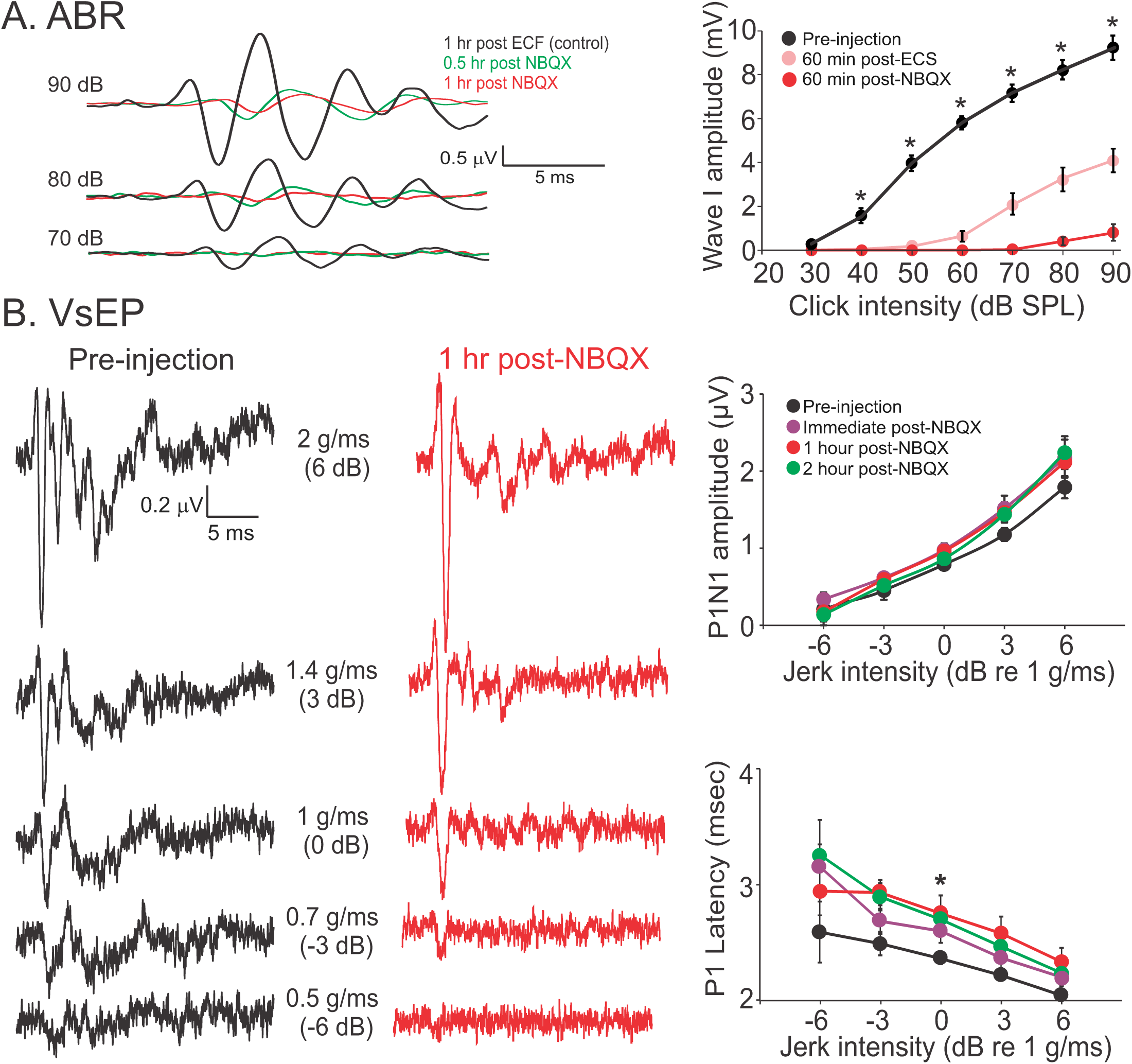
Non-quantal transmission is sufficient for VsEP responses after acute inhibition of glutamate receptors. **A.** Example ABR responses to a click stimulus in a mouse after bilateral IT injections of extracellular solution (ECS) and subsequent IT injections of NBQX solution (20 μM). Responses to three highest click amplitudes are shown. Injection of ECS is used as control, since disruption of the tympanic membrane and filling the canal with fluid result in a decrease in ABR amplitude. The average plot on the right shows pre-injection ABR response (n = 10) compared to after IT injection of ECS (n = 5) or NBQX (n = 5). Based on these results, a decrease of at least 85% in ABR amplitude compared to pre-injection responses for 90 dB click was considered as evidence of penetration of NBQX into the inner ear for the rest of the experiments. Average amplitudes of ABR wave I for 70 – 90 dB stimuli before injection were 7.1 ± 0.4, 8.2 ± 0.4, 9.2 ± 0.5 μV, respectively and 60 min after ECS injection were 2.0 ± 0.5, 3.3 ± 0.5, 4.1 ± 0.5 μV, respectively and 60 min after injection of NBQX were 0 ± 0, 0.3 ± 0.2, 0.8 ± 0.4 μV, respectively. Pre-injection values were larger than post-ECS or post-NBQX values, for intensities of sound above 30 dB (ANOVA, Bonferroni post hoc test, p < 0.001 for all comparisons). Also, post-NBQX values were smaller than post-ECS for sound amplitudes of 70 – 90 dB (ANOVA, Bonferroni posthoc test, p < 0.015 for all comparisons). **B.** Example VsEP responses are shown on the left, for different linear accelerations (i.e., jerks of 0.5 g/ms to 2 g/ms) before (black) and after (red) IT injection of NBQX. There was no change in average amplitudes shown in plots on the right (n = 5, repeated measures ANOVA, p = 0.2). Note, while for 1 g/ms (i.e., 0 dB) stimulus, P1 latencies were significantly larger post-injection compared to pre-injection (repeated measures ANOVA, Bonferroni post hoc test, p = 0.01, 0.04, 0.01 for comparison to immediately, 1 hour, and 2 hours after injection, respectively), the change was in the range of μs and not meaningful from a functional point of view. For the 3 largest stimuli, pre-injection P1N1 amplitudes were 0.8 ± 0.04, 1.2 ± 0.08, 1.8 ± 0.14 μV, immediately after NBQX injection were 0.9 ± 0.04, 1.5 ± 0.09, 2.1 ± 0.16 μV, an hour post injection were 0.9 ± 0.03, 1.4 ± 0.10, 2.2 ± 0.20 μV, and two hours post injection were 0.9 ± 1.0, 1.4 ± 0.35, 2.2 ± 0.25 μV. For the 3 largest stimuli, pre-injection P1 latencies were 2.4 ± 0.05, 2.2 ± 0.02, 2.0 ± 0.01 ms, immediately after NBQX injection were 2.6 ± 0.1, 2.4 ± 0.09, 2.0 ± 0.07 ms, an hour post injection were 2.8± 0.16, 2.6 ± 0.15, 2.3 ± 0.12 ms, and two hours post injection were 2.7 ± 0.11, 2.5 ± 0.10, 2.2 ± 0.11 ms. Data were collected from the same WT mice as in Figure 6.

### Quantal transmission is necessary for normal detection of gravity

Our findings, along with those of other groups indicate that nonquantal transmission via type I hair cells is sufficient to elicit appropriate vestibular afferent and behavioral responses across a broad range of head movement frequencies. The functional role of type II hair cells in amniotes, however, remains unclear. It has been suggested that regular firing afferents provide information about tonic stimuli, such as gravity (Jamali et al., 2019). Bouton-only afferents that have very regular firing receive inputs exclusively from type II HC (Lysakowski et al., 1995) and depend solely on quantal transmission. Therefore, it is possible that afferent responses to tonic stimulation such as gravity would be affected in the absence of peripheral quantal transmission. We used ‘contact righting reflex’ as a measure of sense of gravity (Feather-Schussler and Ferguson, 2016; Grin’kina et al., 2016) before and after IT injection of NBQX in WT mice. When control WT mice (n = 6) were positioned in the supine position between two surfaces, they turned into prone position within 27 ± 6 s (Fig. 8). The same mice after IT injection of NBQX stayed in the supine position and moved around for more than a minute prior to turning to a prone position, with some animals not turning even after 10 min (27 +/- 6 s vs. 437 +/- 90 s, paired t-test, p = 0.009) (Fig. 8). In control experiments, IT injection of extracellular solution did not result in an abnormal contact righting reflex (Before: 28 ± 12 s, After:24 ± 10 s, n = 5, paired t-test, p = 0.4). We also observed abnormal contact righting reflex in vglut3 KO mice (n = 5), with an average of 157 ± 52 s before they turned to a prone position (Fig. 8), which was different from the time for WT mice (t-test, p = 0.03), but not the time required after NBQX injection (t-test, p = 0.07). These results show that peripheral quantal transmission plays a role in detection of gravity (i.e., a constant acceleration stimulus) by the otoliths.

**Figure 8.**
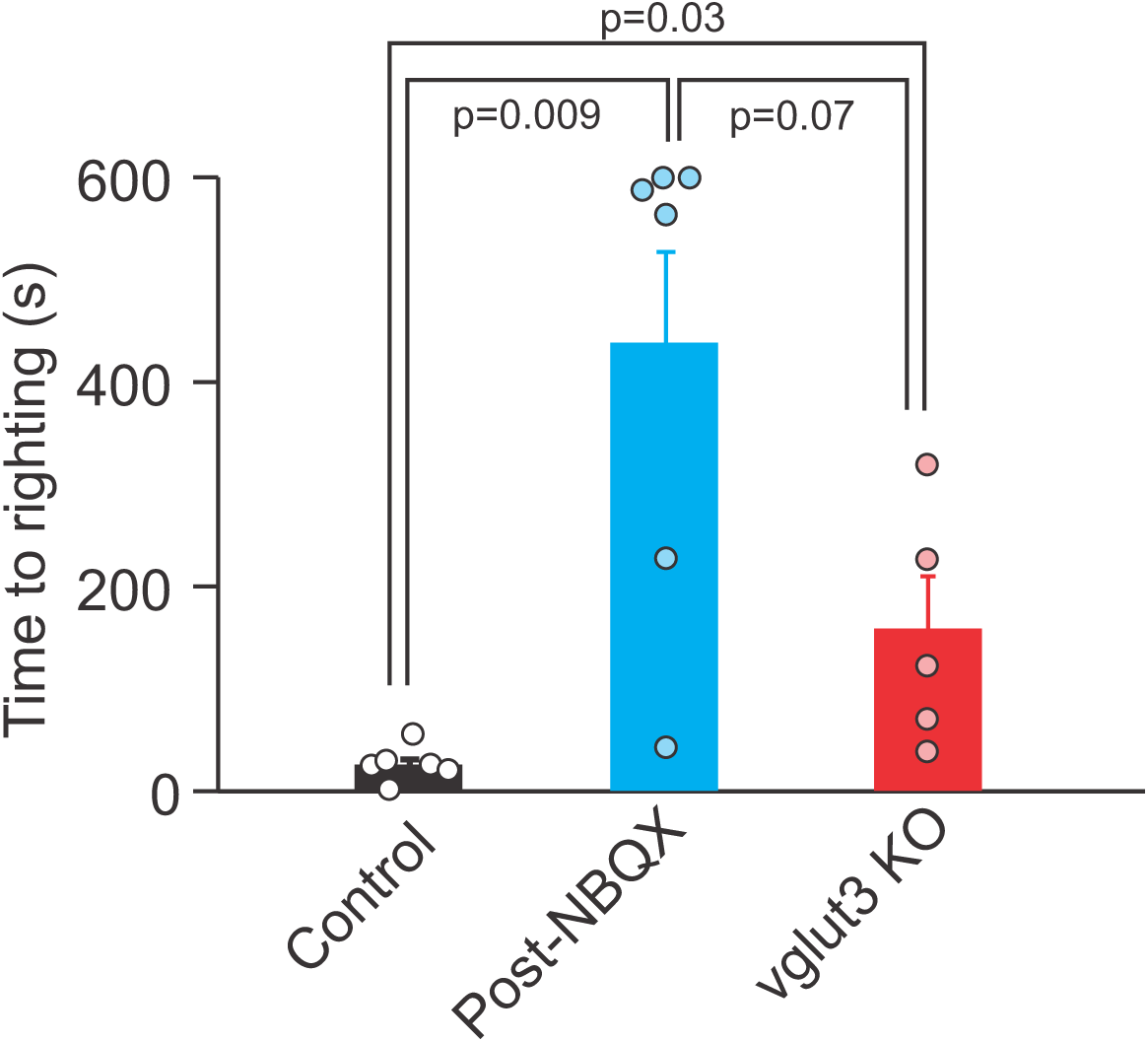
Contact righting reflex is abnormal in the absence of quantal transmission. Mice were in supine position between two flat surfaces and the time for them to flip over to a prone position was measured to quantify each mouse’s ability to sense gravity before and after IT injection of NBQX to block glutamate transmission and compared to values for vglut3 KO mice. Control mice typically flip over in seconds (27 ± 6 s). In the absence of normal quantal transmission after NBQX injections, the animals would stay in the supine position for a long time and even walk around upside down, before they eventually flip over to a prone position (437 ± 90 s, n = 5, paired t-test, p = 0.009). We stopped the experiment at 10 min if the animal did not flip over to the prone position. In addition, for vglut3 KO mice (n = 5, 157 ± 52 s), the time to flip was longer than WT mice (t-test, p = 0.03), but not different from post-NBQX values (t-test, p = 0.07).

## Discussion

We established that NQ transmission between type I HCs and calyx afferent terminals in the vestibular periphery is *sufficient* for providing normal vestibular responses to head movements in the mid to fast range, as evidenced by intact VOR and VsEP results. Although most afferents also receive quantal inputs from type II HCs via bouton and calyceal synapses—and probably from type I HCs as well—we could not assess type I quantal input due to limited optogenetic depolarization. We further show that quantal input is *necessary* for detecting tonic stimuli such as gravity.

### NQ transmission is necessary and sufficient for responses to fast and ultrafast head movements

To evaluate the role of NQ transmission, we used two models with impaired quantal glutamate signaling: vglut3 KO mice, which lack vesicular glutamate release chronically, and mice given intratympanic NBQX to acutely block glutamate receptors. While both models showed reduced or absent ABRs, VsEPs remained intact, consistent with a prior study in guinea pigs (Pastras et al., 2023). This suggests that NQ transmission, with its rapid kinetics, can effectively encode the brief stimuli used in VsEPs. Vglut3 KO mice also exhibited normal VOR responses to head movements up to 1 Hz, suggesting that NQ signaling between type I HCs and calyces can drive VOR, at least over this range of head movements. A recent study has shown that VOR gains are reduced with ototoxic damage selectively in type I HC – calyx synapses in mice (Schenberg et al., 2023), further supporting the role of NQ transmission in VOR generation. Together, these findings suggest that NQ transmission is both necessary and sufficient for normal vestibular signaling during dynamic head movements.

### Peripheral quantal transmission is not necessary for regular resting discharge by afferents

The precise mechanisms underlying variability in vestibular afferent discharge regularity are not fully understood. Anatomically, regular afferents tend to innervate peripheral/extrastriolar zones, while irregular afferents target central/striolar regions (Lysakowski et al., 1995; Brichta and Goldberg, 1996). Calyx terminals in these regions differ in firing properties and potassium channel expression (Lysakowski et al., 2011; Meredith and Rennie, 2015). Furthermore, although most afferents are ‘dimorphic’ and receive inputs from both type I and type II HCs (Fernández et al., 1995a; Lysakowski and Goldberg, 1997, 2008a), those exclusively receiving inputs only from type I HCs in central/striolar zones show highly irregular resting discharge, whereas those receiving inputs only from type II HCs through bouton terminals in the peripheral/extrastriolar region show highly regular resting discharge patterns. For dimorphic afferents, a direct correlation has been shown between the number of bouton terminals (i.e., inputs from type II HCs) and regularity of discharge. Regional differences in sodium channel properties most likely also contribute to these patterns (Liu et al., 2016; Meredith and Rennie, 2018; Baeza-Loya and Eatock, 2024).

Our results suggest that the regularity of discharge is not contingent upon quantal inputs. In vglut3 KO mice, which lack quantal release and thus eliminate the contribution of type II HCs, afferents could still be divided into two groups based on the CV* of 0.15 as the cutoff. We therefore conclude that peripheral quantal transmission and type II HC inputs are not required for regular resting discharge of afferents. Instead, NQ inputs from type I HCs are sufficient for generating various resting firing patterns. This further suggests that the regularity of discharge is mainly dependent on post-synaptic afferent properties between central and peripheral regions of the cristae and maculae, as suggested previously (Goldberg, 2000).

It has been proposed that VOR responses primarily depend on input from regular vestibular afferents, as selective suppression of irregular afferents using galvanic currents did not affect VOR performance (Minor and Goldberg, 1991). Our findings are consistent with this view. The presence of regular afferents in vglut3 KO mice and the normal VOR observed in these mice suggests that type I hair cell input alone can provide sufficient information for VOR generation. This aligns with recent evidence showing reduced VOR gains following damage to calyx-type I hair cell synapses (Schenberg et al., 2023).

We therefore conclude that in the absence of inputs from type II HCs, afferents with regular resting discharges are still present and provide appropriate signals for the generation of normal VOR responses.

### Peripheral quantal transmission is necessary for detection of gravity

In addition to participating in generating the VOR, regular afferents have also been proposed to encode tonic/sustained stimuli, such as changes in head position relative to gravity (Jamali et al., 2019). Here, we showed that despite the presence of regular afferents, the sense of gravity was disrupted in the absence of peripheral glutamatergic transmission. This suggests that quantal transmission in the inner ear is required for detecting tonic stimuli, such as the gravity. This shows that although NQ inputs from type I HCs are sufficient for generating responses by regular afferents during faster head movements during VOR, quantal inputs are required for detection of tonic stimuli.

### Role of type I and type II HCs in transmitting Quantal versus NQ inputs to calyx afferents

Most afferent fibers contact multiple type I and type II HCs. The use of optogenetics enabled the stimulation of a large population of hair cells, providing a more comprehensive view of the combined inputs to afferent fibers than previous studies, which typically activated only single or small groups of nearby HCs via mechano-transduction (Vollrath and Eatock, 2003; Songer and Eatock, 2013; Asai et al., 2018; Ono et al., 2024). While we were unable to drive vesicular release from type I HCs through optogenetic stimulation alone, it is highly possible that type I HCs would not depolarize much under natural conditions due to their low membrane resistance (Rennie et al., 1996, 2004; Rusch and Eatock, 1996; Bao et al., 2003; Holt et al., 2007b; Li et al., 2010; Martin et al., 2024), and therefore the artificial optogenetic stimulation here may be somewhat representative of the relative level of type I and type II HC contributions *in vivo*. However, immunohistochemistry and electron microscopy (Fernández et al., 1995a; Lysakowski and Goldberg, 1997, 2008a; Sadeghi et al., 2014; González-Garrido et al., 2021; Ahmad et al., 2025) confirm the presence of intact synaptic ribbons, indicating that type I HCs are most likely, capable of releasing vesicles onto the inner surface of calyx terminals (Dulon et al., 2009). Previous studies have also reported ribbons in type II HCs along the outer surface of calyx terminals (Lysakowski and Goldberg, 1997, 2008a; Ahmad et al., 2025), suggesting vesicular quantal input from type II HCs onto calyces. The number of ribbons over the calyx outer surface (from type II HC) are fewer compared to its inner surface (from type I HC), but these outer face ribbons are more common in mice compared to chinchillas and squirrel monkeys (Fernández et al., 1995b; Lysakowski and Goldberg, 2008b; Ahmad et al., 2025). The frequent occurrence of synaptic events in our calyx recordings during optogenetic stimulation supports the idea that, in mice, type II HCs may be a predominant source of input to calyx terminals.

We also observed a standing current in calyces with optogenetic stimulation of type II HCs, which was blocked by glutamate receptor antagonist NBQX. This suggests that increased vesicular release during stimulation leads to glutamate accumulation in the synaptic cleft between the type II HC and its neighboring calyx, resulting in a constant inward current that potentially contributes to afferent firing. Although the kinetics of this response was not rigorously evaluated, due to the more gradual accumulation of glutamate it is anticipated to exhibit slower kinetics compared to the rapid NQ transmission.

## Conclusion

During vertebrate evolution from water to land, amniotes (reptiles, birds and mammals) developed type I vestibular hair cells (HCs), paired with unique calyceal afferent nerve fiber terminals that cover the basolateral walls of one or more HCs. Type II HCs retained bouton synapses, similar to those in fish and amphibians. Our results combined with those of previous studies suggest the following: (1) NQ transmission at the specialized synapses between type I HCs and calyx afferent terminals is necessary and sufficient for generating responses to rapid head movements and crucial for normal VOR and VsEP responses; (2) NQ transmission plays a role in encoding head movements by both regular and irregular afferents, facilitating VOR and VsEP responses, respectively; and (3) peripheral quantal transmission is necessary for accurately encoding slower or tonic stimuli, such as detecting gravity.

## Acknowledgements

This work was supported by a National Institute on Deafness and Other Communication Disorders R01 DC012957 to EG and R01 DC019380 to SGS, National Eye Institute R01 EY011027 and National Institutes for Neurological Disorders and Stroke R01 NS132880 to SdL, F31 DC014910 to ZY, the David M Rubenstein Professorship to EG, David M Rubenstein Precision Medicine Center of Excellence in Hearing Fund, and Lloyd B. Minor Center for Vestibular and Skull Base Sciences.

